# Transient matrix remodeling during direct cell-fate reprogramming enables self-assembly of dense neuronal niches

**DOI:** 10.64898/2026.10.27.762125

**Authors:** Kerstin Laurin, Mette Habekost, Fredrik Nilsson, Pinelopi Apostolaki, Emily B. Roloson, Jessica Giacomoni, Andreas Bruzelius, Milan Finn Wesseler, Roger A. Barker, Niels Bent Larsen, Daniella Rylander Ottosson, David J. Mooney, Malin Parmar, Janko Kajtez

**Author notes:** Correspondence: Janko Kajtez.

## Abstract

Native tissues are characterized by high cellular density and continuous reciprocal interactions between cells and the extracellular matrix (ECM), yet recreating these features *in vitro* remains a major challenge. Current three-dimensional (3D) culture systems typically capture either dense cell-cell interactions or a defined ECM environment, but rarely both. Here, we introduce a strategy that exploits direct fibroblast-to-neuron reprogramming to decouple microenvironment formation from final cell identity. Human dermal fibroblasts were embedded in collagen microgels, where their intrinsic matrix-remodeling capacity drove rapid ECM compaction. Concurrently, the contractile fibroblasts were converted into induced neurons (iN), generating densely packed neuronal microcultures within a remodeled collagen matrix. This approach transforms a defining function of the starting cell type into a mechanism for engineering the microenvironment of the target cell type. The resulting iN-microgels combine high cell density, defined ECM composition, and tunable biophysical properties in a scalable microscale format. By coupling matrix remodeling with cell-fate conversion, our platform establishes a general framework for engineering tissue-like microenvironments and reveals how dynamic 3D extracellular contexts can be integrated into direct cell reprogramming.

## Introduction

Native tissues are defined by a dense cellular organization embedded within a complex extracellular matrix (ECM), where cells are simultaneously engaged in extensive cell-cell and cell-ECM interactions. This tightly integrated architecture is not merely structural but instructive, as physical and biochemical cues from the microenvironment are continuously transduced into intracellular signaling pathways that regulate cell identity, function, and fate. Reconstituting these features *in vitro* remains a central challenge, as conventional cell culture formats seldom capture both the high cellular density and the dynamic reciprocal interaction between cells and the surrounding matrix^1^.

Three-dimensional (3D) culture systems have emerged as a promising strategy to better approximate *in vivo* conditions^2–4^. However, existing 3D formats typically recapitulate only selected aspects of tissue organization. Scaffold-free systems promote high cell density and robust cell-cell interactions but lack a defined and tunable ECM component^5, 6^. In contrast, hydrogel-based encapsulation provides a controllable matrix but is typically limited to low cell densities or embedding of pre-formed aggregates leading to highly variable density profile across the construct^7, 8^. In practice, achieving high cell density within hydrogels remains intrinsically challenging because the hydrogel precursor solution occupies most of the formulation volume, thus limiting the fraction available for cells.

Many adherent cell types actively remodel their surrounding ECM through actomyosin-mediated contractile forces that reorganize and compact fibrillar networks^9^. This behavior is well established in collagen gel contraction assays, where mesenchymal cells, including fibroblasts and stromal populations, drive substantial matrix compaction in models of wound healing, fibrosis, and cytotoxicity^10–13^. Similar ECM-remodeling principles have been widely used to generate heart and muscle microtissues inside fibrin hydrogels^14, 15^. Such remodeling can markedly increase local cell densities while simultaneously restructuring the ECM. However, this strategy relies on intrinsically contractile cell types. Neurons have a limited matrix remodeling capacity, making it difficult to establish dense neuronal cultures within an ECM-integrated microenvironment. This raises the question of how similarly dense, ECM-integrated microenvironments can be generated for cell types that lack intrinsic contractility?

We hypothesized that direct fibroblast-to-neuron reprogramming could provide a solution to this challenge by decoupling microenvironment formation from final cell identity. Cells with strong matrix-remodeling capacity could first be used to generate dense, ECM-integrated microenvironments and subsequently converted into induced neurons (iNs), that lack such capabilities, via direct reprogramming, a process that converts one type of mature cell directly into another without first reverting to a pluripotent stem cell state^16^. Since the pioneering experiments demonstrated direct murine fibroblast-to-neuron conversion more than a decade ago^17, 18^, a growing number of protocols have emerged for the generation of targeted neuronal subtypes from adult human dermal fibroblasts (hDFs) using defined cocktails of transcription factors and fate determinants^19^. However, despite extensive optimization of genetic and biochemical cues, many reprogramming approaches are still performed on plastic two-dimensional (2D) substrates. Consequently, the potential of dynamic ECM remodeling and three-dimensional microenvironments to influence neuronal conversion and tissue assembly remains largely unexplored.

Here, we present a 3D platform for direct neuronal reprogramming of adult hDFs inside defined collagen-based microgels, termed iN-microgels (**Fig. 1a**). This approach leverages the intrinsic ability of fibroblasts to remodel and contract collagen matrices, resulting in rapid compaction of the surrounding matrix and the formation of dense, tissue-like microenvironments. While hDFs are contracting the surrounding matrix, lentiviral expression of neuronal reprogramming factors drives their fate conversion into iNs. In this system, one of the fundamental functions of the starting cell type (i.e., fibroblasts) is leveraged to create the microenvironment for the target cell type (i.e., neurons). As a result, iN-microgels enable the scalable generation of densely packed, ECM-integrated neuronal niches, thereby overcoming a fundamental limitation of existing 3D systems by uniting microscale format, high cell density, defined ECM, and tunable biophysical cues within a single platform.

**Fig. 1:**
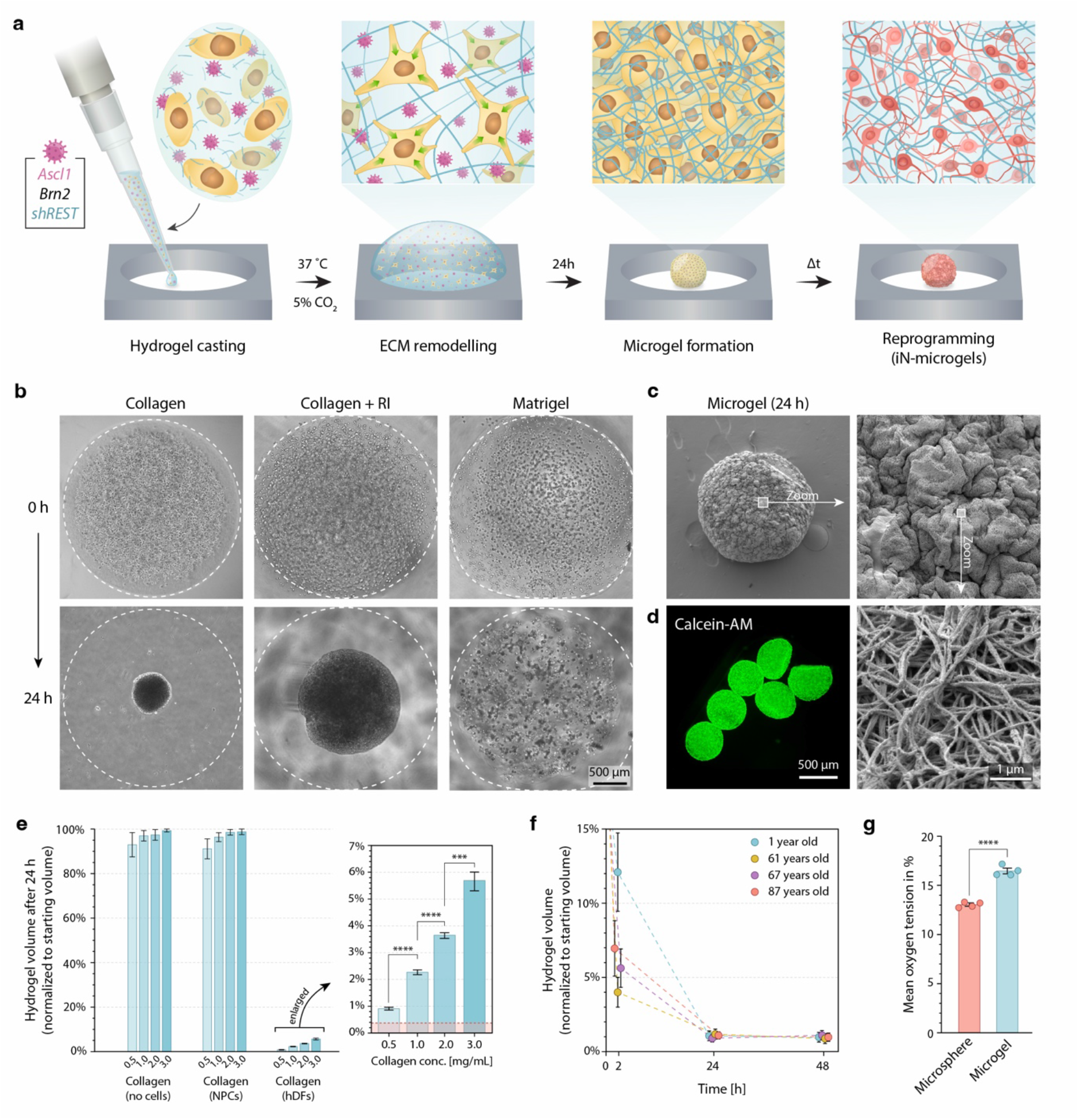
iN-microgel concept and microgel formation. **a** Graphical representation of the proposed concept. **b** Brightfield images demonstrating efficient hDF contraction of collagen hydrogel 24 h after the seeding (left). The process is inhibited by the addition of ROCKi (middle). Contraction is not observed in Matrigel hydrogels (right). **c** Scanning electron microscope images visualizing the remodeled ECM across whole iN-microgel surface (top left) in a form of garified ridges (top right) that are intertwined with collagen fibrils. **d** Visualization of cells using live cell dye (Calcein-AM) within fully contracted microgels. **e** Measurement of contraction after 24 h in collagen hydrogels at four different concentrations (0.5, 1, 2, and 3 mg/mL) and three conditions: plain hydrogels not loaded with cells, hydrogels loaded with neural progenitor cells (NPCs), and hydrogels loaded with hDFs. Enlarged graph demonstrates concentration dependent compaction. Red bar indicates the size of an ECM-free self-assembled hDF microsphere (maximum packing condition). (n = 3) **f** Measurements of hydrogel volume of 0.5 mg/mL collagen hydrogels loaded with hDFs derived from donors of different ages. (n = 6) **g** Oxygen level measurements using oxygen sensitive polystyrene microprobes embedded within microcultures: microsphere^29^ containing no ECM or hDF-compacted collagen microgels. (n = 4, average of 25 probes per microculture construct) All data are mean ± s.d.

## Results

### Efficient compaction of collagen hydrogels by hDFs

The approach proposed here depends on the ability of hDFs to efficiently compact collagen matrix before cell fate switch occurs. While collagen contraction assays are well-established, it is unclear whether hDF exposure to lentiviral vectors and consequential overexpression of neuronal reprogramming factors would interfere with their matrix-remodeling ability. Previous studies indicated that within 48 h of induction of the pioneering factor *Ascl1*, most fibroblasts have already exited the cell cycle and begun remodeling their cytoskeleton^20^. Matrix compaction must therefore be completed within this timeframe. Accordingly, we examined whether hDFs retain this ability following initiation of overexpression of *Ascl1* and *Brn2* together with knockdown of *REST*, delivered by an all-in-one polycistronic lentiviral vector termed pB.pA. ^21, 22^.

Within the first 24 hours, hDFs visibly compacted the cast collagen hydrogel into a spherical microgel (**Fig. 1b left**). This remodeling is known to be driven by actomyosin contractility: fibroblasts attach to surrounding collagen fibers through integrin-based adhesions and pull on them, progressively drawing the network inward. As expected, inhibition of this pathway with a ROCK inhibitor (RI) ^23, 24^ markedly impaired compaction, leading to less compacted gels (**Fig. 1b middle**). Beyond confirming the mechanism, this offers a route to tuning the final microgel size through modulation of contractility. Notably, when hDFs were instead embedded in Matrigel, a matrix composed predominantly of laminin with little fibrillar collagen^25^, the gel compacted only marginally with the cells instead reorganizing into small aggregates dispersed throughout the gel volume (**Fig. 1b right**). This indicates that a fiber-rich network is required for cells to transmit contractile forces across the matrix and compact it. Scanning electron microscopy of the compacted gels revealed a dense network of thick, bundled collagen fibers extending across the entire microgel surface, consistent with physical crosslinking of the fibrils as they are drawn together during compaction (**Fig. 1c**). Live-cell staining by Calcein-AM confirmed that cells remained viable and distributed throughout the microgel volume (**Fig. 1d**).

We reasoned that the final microgel size would be set by a balance of two opposing effects related to collagen concertation, a thug-of-war between fibroblast remodeling capacity and the collagen crosslinking dynamics. Gels with lower collagen concentration contain less fibers to be remodeled which should lead to higher contraction, while gels with higher collagen concentration offer more fibers that should require longer time to be remodeled and oppose the change through physical crosslinking of collagen. To test this, we cast hDF-laden gels at four collagen concentrations (0.5, 1, 2 and 3 mg/mL) and compared their volume 24 h after casting to the volume at casting.

All conditions contracted substantially, reaching 1–6% of their starting volume depending on collagen concentration (**Fig. 1e**). Compaction scaled inversely with concentration: 0.5 mg/mL gels contracted the most, to 0.9 ± 0.1% of the initial volume, followed by 1 mg/mL (2.2 ± 0.1%), 2 mg/mL (3.6 ± 0.1%) and 3 mg/mL (5.6 ± 0.3%). Cell-free collagen gels retained 99.5 ± 0.15% of their volume over the same period indicating minor hydrogel-intrinsic reduction in volume. Gels seeded with neural progenitor cells (NPCs), cells with limited ECM-remodeling capacity, in place of hDFs retained 98.9 ± 0.5%, confirming that compaction requires hDFs specifically rather than the mere presence of embedded cells. Finally, we asked whether donor age affects this behavior by comparing hDF lines from donors aged 1 to 87 years. Compaction dynamics differed between lines over the first 2 hours, but these differences resolved with time and were absent at both 24 and 48 h indicating no age-related impairment of contraction efficiency of hDFs (**Fig. 1f**). Importantly, virus transduction did not affect compaction efficiency (**Fig. S1a**)

While the collagen-contraction approach yields high-cell-density constructs, their final size remains larger than that of hDF microspheres formed in low-attachment plates purely through self-organization, without a hydrogel. Thus, the remodeled collagen network should therefore not only support a high cell density but also improve oxygen tension within the construct. To test this, we measured mean oxygen tension in the microgels using oxygen sensitive microbeads and compared it with self-aggregated spheroids (microspheres) lacking ECM. Oxygenation was indeed higher in the microgels, with a mean oxygen tension of 16.6 ± 0.5% compared to 13.1 ± 0.3% in the microspheres (n = 4, average of 25 oxygen probes per replicate) (**Fig. 1g** and **S1b**). These measurements were further corroborated by numerical modeling of oxygen distribution indicating 16.3% and 13.5% oxygen in microgels and microspheres respectively (**Fig S2**). Together, these results show that hDFs remodel collagen fibers even in the presence of reprogramming factors, generating compact microgels that combine a high cell density with a supporting ECM, establishing a structured microenvironment for the iNs.

### High-cell density microgels support direct neuronal reprogramming

With the high-cell-density format established in the presence of reprogramming factors, we next asked whether these factors would drive fibroblast-to-neuron conversion within the formed microgels. Using the pB.pA lentiviral vector described above, we analyzed reprogramming outcomes 28 days post-transduction. Immunocytochemistry revealed that cells acquired a neuronal morphology, with small cell bodies and long, thin projections, positive for the neuronal markers TAU, TUBB3 and MAP2, confirming successful conversion of hDFs into iNs and demonstrating for the first time the generation of iN-microgels (**Fig. 2a**). Scanning electron microscopy further confirmed neuronal morphology and revealed direct contact between the neurons and the surrounding collagen network (**Fig. 2b**). Importantly, confocal fluorescence microscopy of optically cleared iN-microgels showed that reprogramming occurred throughout the volume of the cast hydrogel, not only at its surface (**Fig. 2c**).

**Fig. 2:**
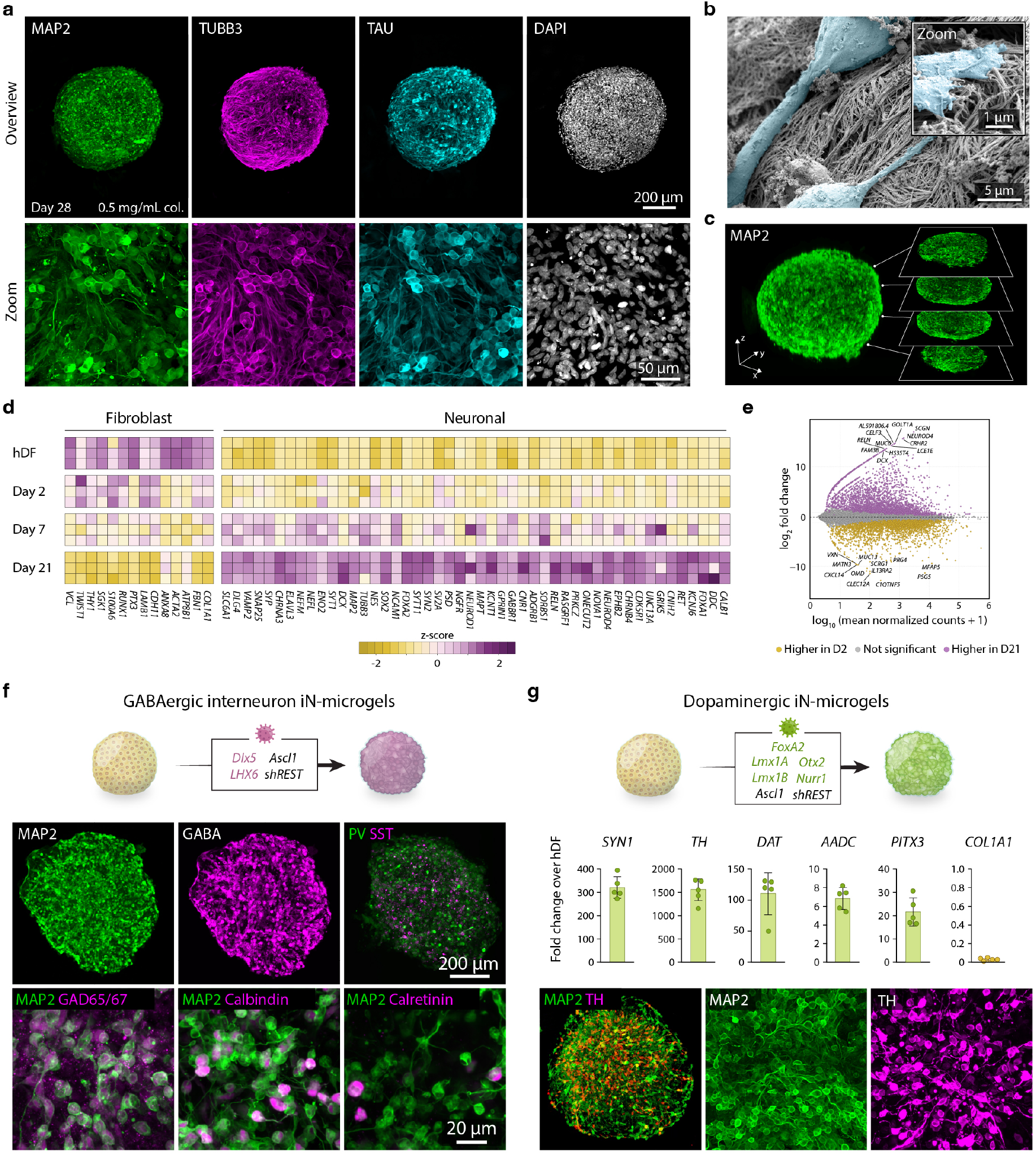
Direct neuronal reprogramming inside compacted hydrogel microcultures. **a** Immunocytochemistry of 0.5 mg/mL iN-microgels after 28 days of reprogramming showing expression of neuronal markers MAP2, TUBB3 and TAU. Overview of whole microgel on top with a zoomed in image on the bottom to show neuronal morphology. **b** Scanning electron image of reprogrammed neurons in connection with the collagen fibers of the iN-microgel. **c** Overview image of iN-microgel showing expression of MAP2 throughout the whole microgel. **d** Heatmap representation of selected fibroblast-associated and neuronal genes. Values were log transformed and scaled by the mean expression per gene across samples. **e** MA-plot of differentially expressed (DE) genes (P adj > 0.05) between iN-microgels at day 2 and day 21. Upregulated DE genes with log_2_ fold change > 1 in day 21 samples are shown in purple and downregulated genes with log_2_ fold change < −1 in yellow. **f** GABAergic interneuron iN-microgels expressing neuronal and interneuron specific markers shown with immunocytochemistry. **g** Dopaminergic iN-microgels, RT-qPCR analysis of *SYN1*, *TH*, *DAT*, *AADC*, *PITX3* and *COL1A1* and immunocytochemistry for MAP2 and TH. (n = 5) All data are mean ± s.d.

Bulk RNA sequencing of hDFs and iN-microgels at days 2, 7 and 21 revealed a progressive transition in transcriptional identity across a panel of 14 fibroblast-associated and 47 neuronal genes (**Fig. 2d**). Fibroblast genes including *COL1A1*, *ACTA2*, *TWIST1*, *THY1* and *FBN1* were gradually downregulated, while neuronal genes spanning cytoskeletal (*MAP2*, *TUBB3*, *MAPT*), neurodevelopmental (*DCX*, *NEUROD1*) and synaptic (*SNAP25*, *SYP*, *DLG4*, *SYN2*) categories were progressively upregulated, with most of the change occurring between day 7 and day 21. These results were further confirmed by RT-qPCR of samples from day 7, 14, 21 and 28 which also captured the steady increase of neuronal genes *MAPT*, *MAP2*, *SYN1,* and *SYP* over time while the fibroblast gene *COL1A1* was already strongly reduced at day 7, decreased further at day 14, and remained at that level at later timepoints (**Fig. S3**). To quantify the extent of this transition, we performed differential expression analysis of bulk RNA sequencing data comparing samples 2 days after casting with day 21. Over 5,000 genes were differentially expressed (adjusted P < 0.05, absolute log2 fold change > 1), with 2,765 upregulated and 2,476 downregulated, reflecting a reorganization of the transcriptome consistent with a major change in cellular identity (**Fig. 2e**).

The pB.pA reprogramming strategy induces neuronal conversion without explicitly directing cells towards a specific neuronal subtype but for disease modeling and regenerative applications this becomes central. To assess the adaptability of our approach to neuronal subtype specification, we incorporated previously published lentiviral cocktails to generate induced GABAergic^26^ and dopaminergic neurons^27^ and applied these into our iN-microgel protocol. To generate induced GABAergic interneurons *Ascl1*, *DLX5* and *LHX6* were overexpressed with *REST* knockdown. The resulting neurons reprogrammed within the iN-microgels expressed neuronal marker MAP2 and interneuron markers gamma-aminobutyric acid (GABA) and GAD65/67 (**Fig. 2f**). The cells also expressed interneuron subtype markers such as calbindin, calretinin, somatostatin (SST), and even parvalbumin (PV), which has not been shown with this TF combination^26, 28^. To generate induced dopaminergic neurons (iDANs) *Ascl1*, *Lmx1A*, *Lmx1B*, *Nurr1*, *FoxA2*, and *Otx2* were overexpressed in combination with *REST* knockdown in the iN-microgels. Gene expression analysis of iDAN-microgels showed upregulation of synaptic gene *SYN1* and dopaminergic genes *TH*, *DAT*, *AADC*, *PITX3* with fibroblast gene *COL1A1* being downregulated (**Fig. 2g**). This was also reflected on protein level where they expressed MAP2 and dopamine marker tyrosine hydroxylase (TH).

Bulk hydrogels are poorly suited to intracerebral transplantation, as their gelation dynamics are hard to control during injection and risk clogging the cannula, while pre-crosslinked bulk hydrogel constructs are too large to deliver through a needle. The microgel format avoids both constraints, and we therefore developed a strategy for direct intracerebral transplantation of pre-formed iN-microgels. iN-microgels transplanted into the striatum of immune deficient rats were successfully engrafted, with surviving neurons projecting into the host tissue 4 weeks post-transplantation (**Fig. S4a**). Notably, stacking of individual microgels left cavities within the graft, which can be resolved by mild enzymatic collagen pretreatment prior to injection (**Fig. S4b**). iN-microgels thus enable direct transplantation of ECM-embedded iNs and open new opportunities for reprogramming strategies in transplantation studies.

### ECM support promotes neuronal maturation during reprogramming

We next asked whether the exogenous ECM itself, rather than the 3D format alone, influences reprogramming outcome. We therefore compared iN-microgels with iN-microspheres, generated by self-assembly of hDFs in microwells according to our published approach^29^ (**Fig. 3a**). Both formats were collected for bulk RNA sequencing. Principal component analysis separated hDFs from all reprogrammed samples and showed that iN-microgels and iN-microspheres overlapped at early timepoints (day 2 and 7) but diverged by day 21 (**Fig. 3b**). Differential expression analysis at day 21 identified 1,108 genes upregulated in microgels and 1,227 in microspheres (adjusted P < 0.05, absolute log2 fold change > 1) (**Fig. 3c**). Gene ontology (GO) analysis showed that microspheres were enriched for terms related to ECM and extracellular structure organization and to chemotaxis, whereas microgels were enriched for synaptic transmission, regulation of postsynaptic membrane potential, and regulation of neurogenesis (**Fig. 3d**), consistent with microspheres continuing to build their own matrix while microgels, already provided with one, progressed further toward a neuronal program. Marker-level comparison across fibroblast, neural progenitor, pan-neuronal, synaptic maturation, and neuronal subtype gene sets supported this, with microgels showing both more complete downregulation of fibroblast genes and higher expression of synaptic maturation genes (**Fig. 3e**). Together, these data indicate that the presence of ECM influences reprogramming outcome, most notably for genes involved in synaptic transmission and maturation.

**Fig. 3:**
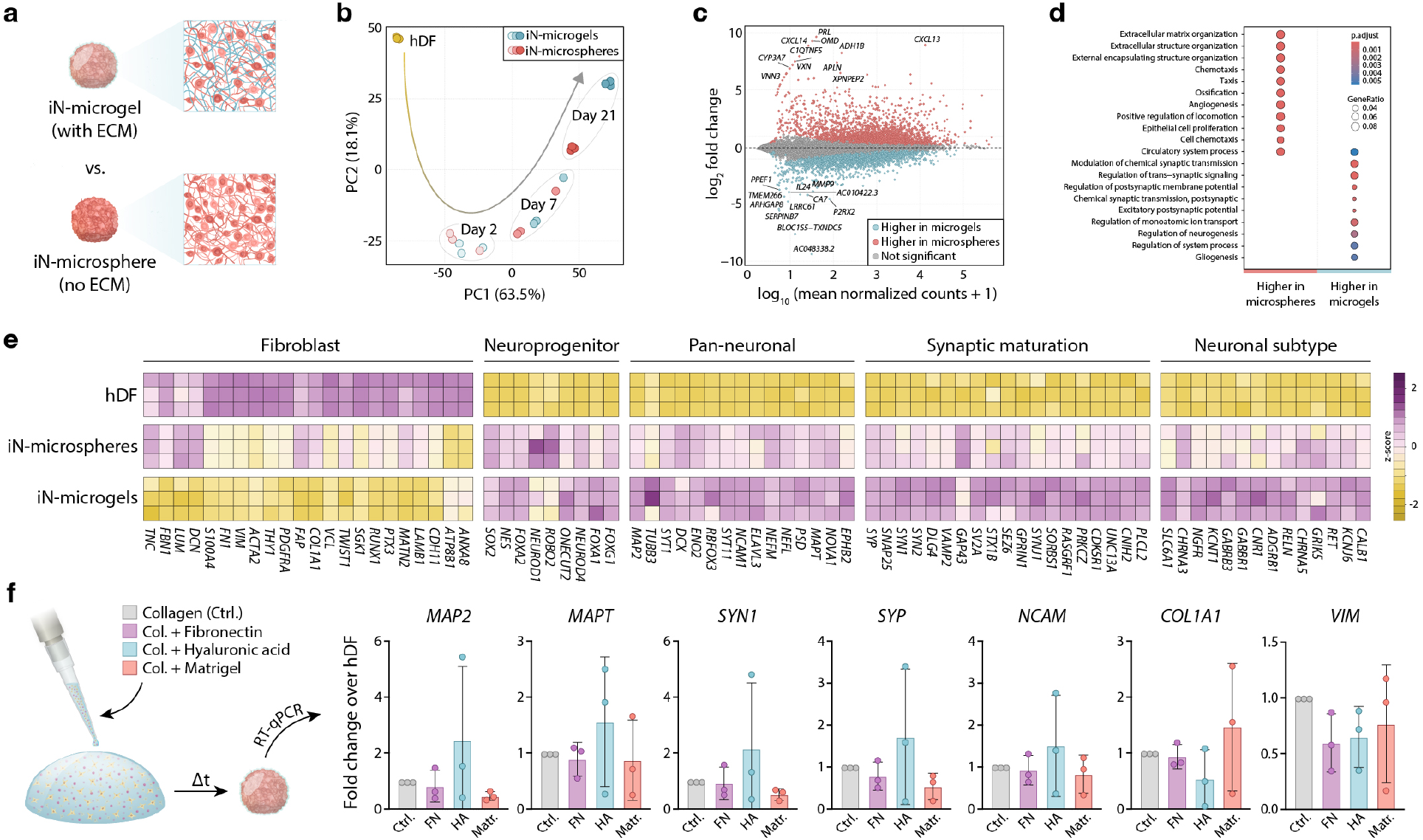
The presence of exogenous ECM enhances direct neuronal reprogramming. **a** Schematic overview of the two compared conditions: 0.5 mg/mL iN microgels and iN-microspheres self-assembled in ultra-low attachment microwells. **b** Principal component analysis (PCA) of transcriptional profiles from hDFs and iN-microgels and iN-microspheres at 2, 7 and 21 days. **c** MA plot of DE genes (P adj < 0.05) between iN-microgels and iN-microspheres at day 21. Upregulated DE genes with log_2_ fold > 1 for iN-microspheres shown in red; upregulated DE genes in iN-microgels with log_2_ fold < −1 shown in blue. **d** Gene ontology (GO) analysis showing GO terms highly expressed in iN-microspheres versus iN-microgels. **e** Heatmap representation of gene expression levels of selected groups of genes in hDF and day 21 iN-microspheres and iN-microgels. Values were log transformed and scaled by the mean expression of each gene across samples. **f** RT-qPCR comparison of selected fibroblast and neuronal genes for iN-microgels with additional ECM components added to iN-microgel mixture. Data shown as fold change over hDF and was normalized to control iN-microgels without added ECM components for each batch. (n = 3, for all statistical comparisons P > 0.05) All data are mean ± s.d.

Collagen I is an abundant ECM component and its fibrillar architecture is what makes compaction possible, but it is not the dominant matrix of the brain. We therefore asked whether supplementing the collagen network with additional ECM components could further support reprogramming, with collagen retained as the structural backbone and the added components acting as adjuvants. Collagen hydrogels were combined with fibronectin (FN), hyaluronic acid (HA), or Matrigel, and gene expression was assessed by RT-qPCR after 21 days of reprogramming and compared with collagen-only iN-microgels (**Fig. 3f**). No condition differed significantly from control, and batch-to-batch variability was high. Nevertheless, collagen combined with HA, a major component of brain ECM, showed a trend toward increased expression of the neuronal genes *MAP2*, *MAPT*, *SYN1*, and *SYP*, together with reduced expression of the fibroblast gene *COL1A1* (**Fig. 3f**). This suggests that the compaction-competent collagen scaffold can accommodate additional matrix components, and that tuning this composition represents a tractable route to improving reprogramming within the microgel format.

### Mechanical properties influence cell fate conversion

We demonstrated that the compaction of iN-microgels was tunable by adjusting collagen concentration. Apart from affecting the final microgel size, we hypothesized that change in collagen concentration would also affect the microgel stiffness. The mechanics of acellular and cell-laden bulk collagen gels are well characterized^30, 31^, and cell-driven compaction is known to stiffen such matrices^32^, but how these properties evolve in the densely compacted microgel format remains unknown. To assess this, we performed nanoindentation on iN-microgels at four collagen concentrations, 3 days after compaction. Young’s modulus scaled with collagen concentration, increasing from 0.89 ± 0.73 kPa (0.5 mg/mL), 1.54 ± 0.79 kPa (1 mg/mL), 2.41 ± 1.36 kPa (2 mg/mL) to 3.87 ± 2.12 kPa (3 mg/mL) (**Fig. 4a**), confirming that collagen concentration sets both final microgel size and stiffness.

**Fig. 4:**
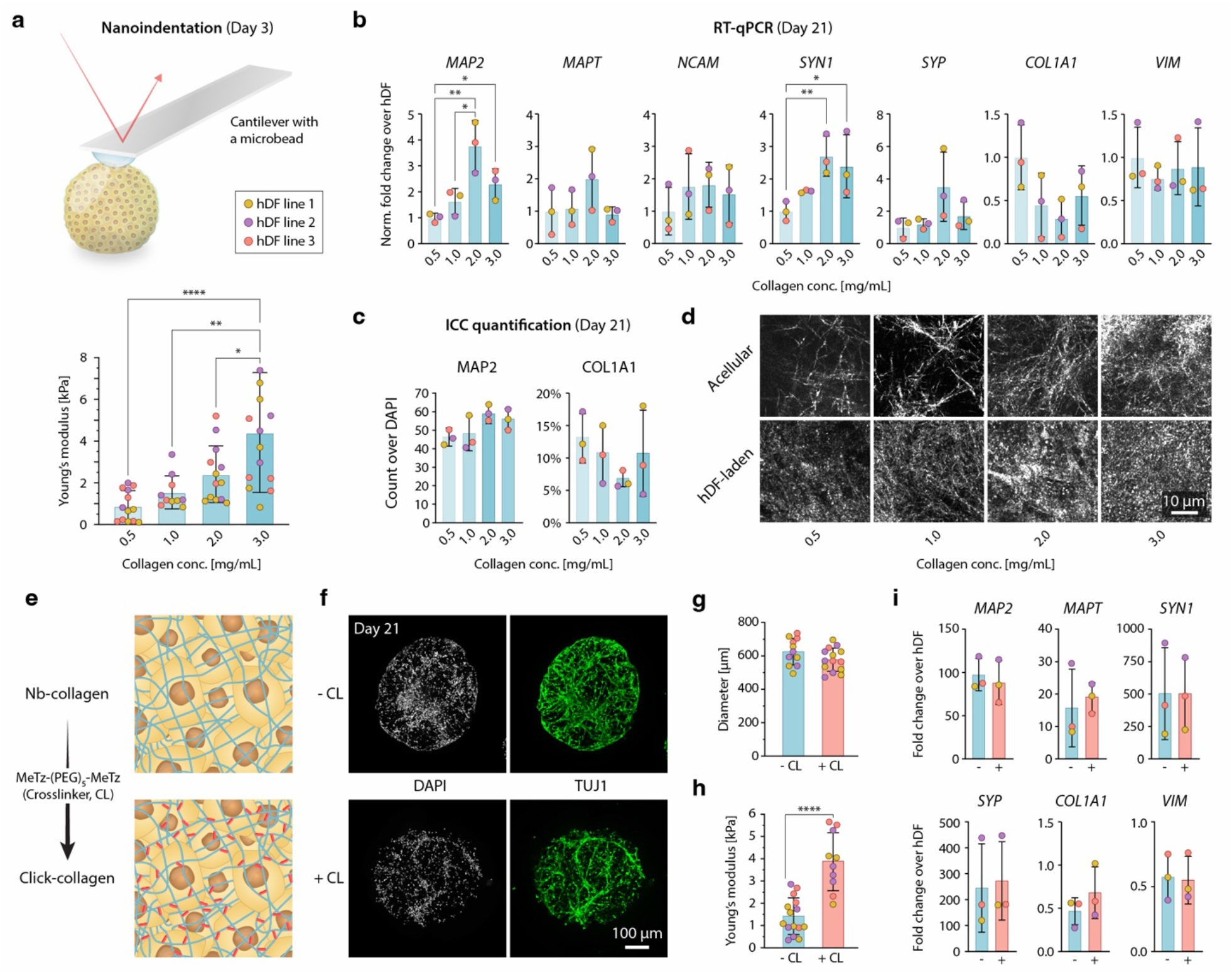
Collagen matrix composition influences direct neuronal reprogramming. **a** Elastic stiffness of collagen microgels measured by nanoindentation after 72 h. Young’s modulus increased with increasing collagen concentration. Each color represents an independent hDF cell line. **b** RT-qPCR analysis of selected fibroblast and neuronal genes in iN-microgels generated at different collagen concentrations. Expression values were normalized to the mean expression at 0.5 mg/mL collagen. Three independent experiments were performed for each cell line. **c** Quantification of neuronal (MAP2) and fibroblast (COL1A1) protein expression from immunofluorescence images of iN-microgels formed at different collagen concentrations. **d** Reflectance microscopy of acellular collagen hydrogels and hDF-laden microgels after contraction, revealing differences in the fibrous collagen network across starting collagen concentrations. **e** Schematic of click-collagen formation. Collagen was functionalized with norbornene (Nb) groups, enabling covalent crosslinking through a divalent methyltetrazine-based linker. **f** Immunocytochemistry for nuclear (DAPI) and neuronal (TUJ1) markers of nb-collagen iN-microgels at day 21 post transduction, without (top, −CL) and with (bottom, +CL) the crosslinker added. **g** Dimeter measurements of nb-collagen iN-microgels, with and without the crosslinker added. **h** Nanoindentation measurements of Young’s modulus of 2 mg/mL collagen microgels with and without covalent crosslinking. **i** RT-qPCR analysis of the selected neuronal (*MAP2*, *MAPT*, *SYN1*, *SYP*) and fibroblast (*COL1A1*, *VIM*) genes in nb-collagen iN-microgels with and without collagen crosslinking. All data are mean ± s.d.

We next asked whether this translates into differences in reprogramming outcome. Microgels of all four concentrations were reprogrammed for 21 days and assessed by RT-qPCR and ICC. Reprogramming was most efficient at an intermediate concentration rather than at the stiffest condition. At 2 mg/mL, the neuronal genes *MAP2* and *SYN1* were significantly upregulated relative to 0.5 mg/mL (**Fig. 4b**), and *MAPT* and *SYP* showed the same trend, together with the lowest levels of the fibroblast gene *COL1A1*, although these differences did not reach significance. This was further confirmed by the ICC where the highest percentage of cells positive for neuronal marker MAP2, 59.3 ± 4.2%, and lowest percentage of cells positive for fibroblast marker COL1A1, 6.4 ± 1.3% was seen in the 2 mg/mL iN-microgels (**Fig. 4c**). Quantifications for the other concentrations showed a lower percentage of MAP2^+^ cells, 45.8 ± 4.1% (0.5 mg/mL), 47.3 ± 9.4% (1 mg/mL), and 54.9 ± 4.9% (3 mg/mL), with a higher proportion of cells being COL1A1^+^, 12.6 ± 3.8 % (0.5 mg/mL), 10.2 ± 4.4 % (1 mg/mL), and 10.1 ± 6.8 % (3 mg/mL). Interestingly, we observed that increasing the collagen concentration during GABAergic conversion significantly and markedly increased expression of key interneuron genes, including *SYN1*, *PV*, *NPY*, *CCK*, *VGAT*, and *GAD2*, while *SST*, *CALB1*, *CALB2*, and *VIP* trended upward without reaching significance, and the fibroblast marker *COL1A1* was further reduced (**Fig. S5**).

To relate these outcomes to network architecture, we imaged the collagen by reflectance microscopy. In acellular gels, fiber density increased with concentration as expected: individual, well-separated fibers were resolved at 0.5 mg/mL, progressively bundled at intermediate concentrations, and at 3 mg/mL formed a mesh too dense to resolve individual fibers (**Fig. 4d**). In hDF-laden microgels the network was considerably denser than in acellular gels of matched concentration, with individual fibers no longer resolvable at any concentration and only larger bundles visible at 0.5 and 1 mg/mL. Cell-driven compaction thus densifies the network well beyond what collagen concentration alone achieves, with a 0.5 mg/mL cell-laden microgel approaching the apparent density of a far more concentrated acellular gel. In the higher concentrations, 2 and 3 mg/mL, images are highly saturated, and it is no longer possible to differentiate individual fibers and structures.

Varying collagen concentration altered stiffness, but it simultaneously altered final microgel size and network density, leaving the contribution of stiffness alone unresolved. We therefore sought a way to modify mechanical properties while keeping the construct otherwise unchanged. Crosslinking the collagen network after compaction offers this, since the microgel has already reached its final size and cell density by 24 h and subsequent crosslinking acts only on the mechanics. For this, we utilized a functionalized norbornene collagen (Nb-collagen) that covalently crosslinks with a divalent methyltetrazine-based linker (**Fig. 4e**). We selected 2 mg/mL for the Nb-collagen microgels, the concentration most beneficial for reprogramming in the previous experiment, and added crosslinker to half the microgels 24 h after casting, once compaction was complete. Nanoindentation at day 3 showed a significant increase in Young’s modulus in crosslinked compared with non-crosslinked microgels, confirming that crosslinking stiffens the construct without altering its size (**Fig. 4f**). We then assessed the effect of crosslinking on the reprogramming outcomes by RT-qPCR on day 21 (**Fig. 4g**). No significant differences in gene expression of neuronal genes (*MAP2*, *MAPT*, *SYN1* and *SYP*) nor fibroblast-associated genes (*COL1A1* and *VIM*) between the crosslinked and non-crosslinked were observed indicating that increased stiffness, at constant microgel size and cell density, does not affect reprogramming outcome in this system.

### Programed shape transformation enables defined 3D geometries

During tissue development, complex 3D architectures emerge through the coordinated generation and redistribution of mechanical forces. Differential contraction and compaction of the ECM can generate local stresses, driving changes in curvature that produce folding and other shape transformations. These mechanically regulated processes enable relatively simple initial geometries to be transformed into highly structured tissue architectures essential to tissue function. Such principles of mechanically driven shape transformation have also been harnessed *in vitro* to generate programmable tissue contractions and geometries^33–35^. Here we exploit these principles to generate neural construct with defined geometries, jointly programming both cell identity and shape.

First, we employed extrusion bioprinting (**Fig. 5a**) to precisely control hydrogel deposition onto a solid substrate. Whereas hydrogel constructs had previously been manually pipetted, extrusion bioprinting enabled automated and spatially controlled deposition. We printed continuous hydrogel filaments with a diameter of 2 mm, and a length of 40 mm. Subsequent hDF-mediated compaction reduced the construct to a continuous cellular structure approximately 20 mm in length and 0.8 mm in diameter. hDFs within the printed constructs successfully underwent direct reprogramming into iNs expressing TUBB3, resulting in continuous, high-cell-density neuronal tissue.

**Fig. 5:**
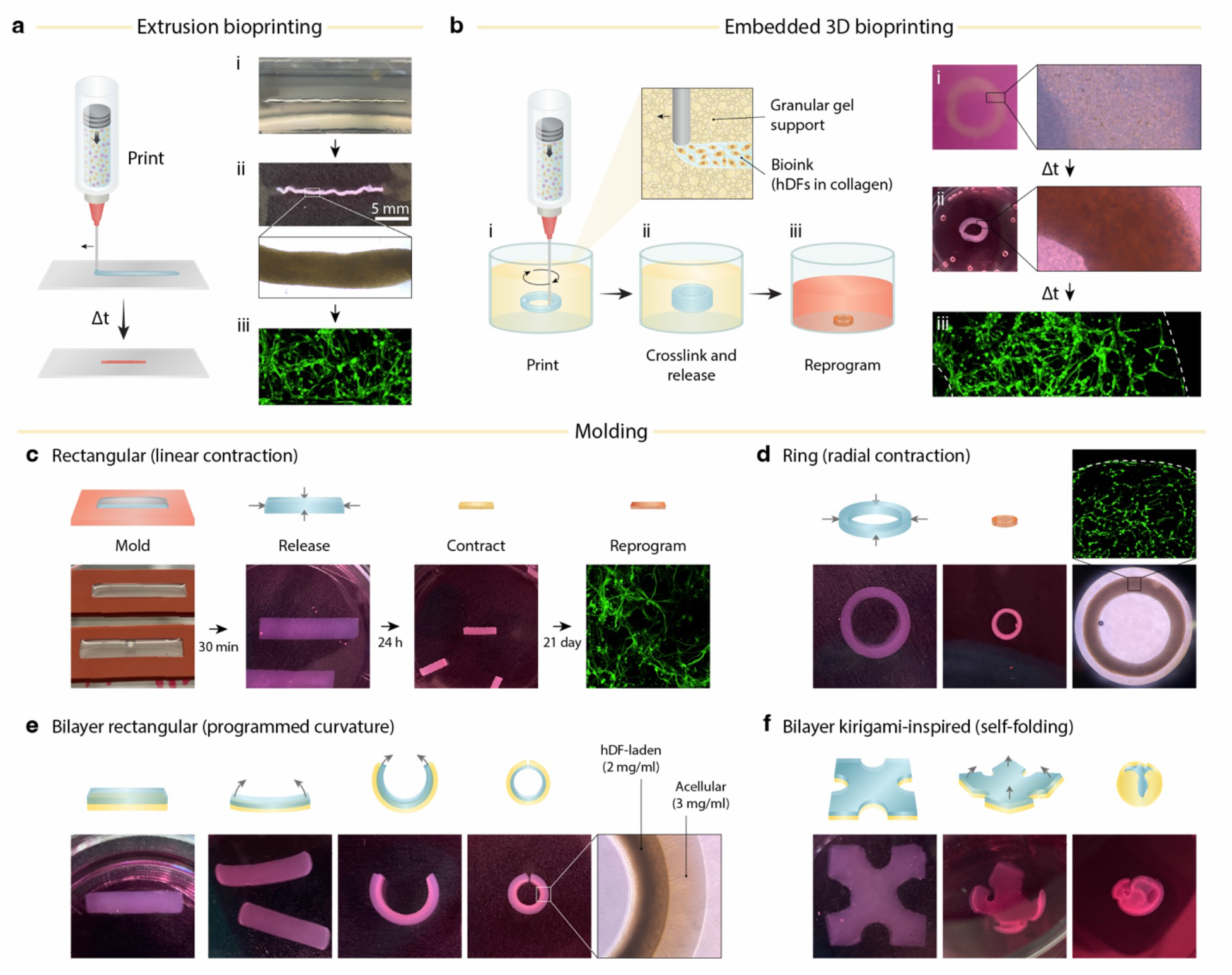
Biofabrication approaches for programmed shape transformation. **a** Extrusion bioprinting of hDF- and lentivirus-laden collagen hydrogel (2 mg/mL) through a syringe needle to generate continuous linear constructs 30 mm in length (i). hDF-mediated matrix compaction produces a dense elongated filamentous construct (ii), within which lentiviral expression of neuronal transcription factors drives direct neuronal reprogramming (iii). **b** Embedded bioprinting of hDF-laden collagen hydrogel within an alginate granular support to generate a cylindrical shell construct (i). Following in situ collagen crosslinking and partial contraction, the construct is released from the support and transferred to culture medium containing lentiviral vectors (ii), resulting in dense neuronal structure (iii). **c** Casting of hDF- and lentivirus-laden collagen hydrogel within a rectangular silicone mold generates defined planar geometries. hDF-mediated linear contraction compacts the construct within 24 h, followed by neuronal reprogramming during culture (21 days). **d** Radial contraction of a ring-shaped construct generates a compact circular geometry with dense neuronal tissue. **e** Bilayer casting of an acellular collagen layer and an hDF-laden collagen layer generates a contractile–non-contractile composite. Contraction of the upper cellular layer induces differential strain and curvature toward the contractile layer, producing a defined circular geometry. **f** Application of the same bilayer architecture to a kirigami-inspired geometry with lateral semicircular cut-outs directs contraction-induced folding into a compact, sphere-like construct.

Conventional extrusion bioprinting is limited in its ability to fabricate complex 3D geometries from soft, low-viscosity pre-gel solutions, as deposited structures can deform or collapse before gelation. To overcome this limitation, we employed embedded 3D bioprinting within a granular support medium (**Fig. 5b**). Granular gel supports enable precise 3D patterning of soft and liquid biomaterials while providing a mechanically compliant environment that permits subsequent cell-mediated compaction of the printed construct^34, 36, 37^. We continuously printed a cylindrical construct (8 mm diameter, 5 mm height) from a collagen–hDF bioink within an alginate granular gel. Following printing, the collagen matrix was crosslinked in situ, after which the construct was allowed to undergo partial hDF-mediated compaction within the support. The compacted construct was then released from the granular matrix and exposed to viral vectors to initiate direct neuronal reprogramming. This process yielded a monolithic, ring-like construct with high cellular density that was subsequently reprogrammed into a continuous, high-cell-density neuronal tissue.

Molding provides a simple approach to generate defined geometries and explore programmable tissue folding under controlled geometric constraints. We therefore fabricated custom silicone molds to pattern collagen-hDF constructs containing the viral vectors for direct neuronal reprogramming. A rectangular construct (6 × 3 mm) was cast, crosslinked, and released into culture medium, where hDF-mediated contraction reduced it to approximately 1 × 3 mm while preserving its rectangular geometry (**Fig. 5c**). A similar approach using a ring-shaped mold generated a compact circular construct through radial contraction (**Fig 5d**).

To introduce directional control over shape transformation, we next fabricated bilayer constructs comprising an acellular collagen layer (3 mg/ml) and an hDF-containing collagen layer (2 mg/ml). Contraction of the cellular layer generated differential strain across the bilayer, producing anisotropic folding and curvature toward the contracting layer and enabling defined circular geometries (**Fig. 5e**). Finally, we applied the same principle to a kirigami-inspired bilayer with a square geometry containing semicircular cut-outs on each side. Contraction of the hDF-containing layer drove self-folding of the initially planar construct into a compact, sphere-like geometry (**Fig. 5f**). Together, these configurations demonstrate programmable control of tissue geometry through spatially constrained cellular contraction.

## Discussion

High cell density and a defined ECM are both hallmarks of tissue, yet they are rarely achieved together *in vitro*. This is particularly challenging for cell types such as neurons, which have limited capacity to remodel their extracellular environment. Here, we address this challenge by using direct fibroblast-to-neuron reprogramming to decouple tissue assembly from cell identity, generating target cells directly within a pre-engineered, cell-dense ECM. This approach exploits fibroblasts not only as starting cells for cell-fate reprogramming, but also as active architects of the extracellular environment, allowing the cell that remodels the matrix to ultimately become the cell that inhabits it.

The versatility of the platform is here demonstrated by its compatibility with three transcription factor-based reprogramming strategies generating distinct neuronal populations, including dopaminergic^27^ and GABAergic neurons^26^. Because the biomaterial environment is established independently of the specific reprogramming factors, the platform could be readily combined with other transcription factor-, mRNA-, or small-molecule-based approaches to generate additional CNS and peripheral neuronal populations, including motor^38^ and sensory^39, 40^ neurons, as well as glial populations such as astrocytes^41, 42^ and oligodendroglia^41^. The same principle could extend to other non-neural lineages, including hepatocytes^43^ and osteoblasts^44^. This flexibility creates an opportunity to tailor the extracellular environment to the requirements of each target cell type while maintain high-cell density context leveraged for disease modeling or regenerative medicine applications.

Major efforts in the field have gone into optimizing reprogramming cocktails and the cell-intrinsic mechanisms limiting conversion, while extrinsic biophysical cues from the ECM remain underutilized. We show that both the presence and the amount of ECM modulate reprogramming outcomes, and our data further indicate that ECM components beyond collagen would be beneficial. Our findings therefore add to the evidence that biophysical cues shape direct neuronal conversion, previously demonstrated on 2D substrates for topography^45–47^ and stiffness^48–50^, and in conventional low-density 3D hydrogel cultures^24, 51, 52^. While our system uniquely promotes high-cell-density hydrogel constructs, a collagen-only matrix makes it difficult to decouple viscoelastic properties, adhesion motif density, and crosslinking extent, since all scale with collagen concentration and with the degree of compaction. Separating them will require fibrillar networks with independently tunable properties that fibroblasts can remodel. Synthetic fibrillar gels^34, 53, 54^ and functionalized natural networks meet this requirement, with three strategies envisioned to advance our system: grafting bioactive peptides, additional adhesion motifs and protease-cleavable sites onto the fibers themselves^55–57^; adding a second, interpenetrating network to set stiffness and stress relaxation independently of the fibrillar architecture^58–60^; and applying orthogonal crosslinking to tune the same viscoelastic properties once compaction is complete^61, 62^, as we show here with click-collagen. Realizing these strategies will also require resolving viscoelasticity within the microcultures themselves, which nanoindentation captures only partially. Capillary micromechanics^63^, Brillouin microscopy^64^, and magnetic microrheology^65^ could provide the complementary readouts needed to link defined matrix properties to reprogramming outcomes.

Here we used printing, molding and bilayer geometries to direct contraction into defined shapes, demonstrating engineered morphogenesis in an origami-like manner. All of these, however, impose force fields externally through set boundaries defined by bioprinting or casting. Programmable shape morphing of more complex architectures could be achieved by encoding them internally by, for example, spatially patterning cell-laden and acellular regions within a single construct^66^, combining cellular microcultures with inert or degradable microgels into granular composites^35^, and defining anchoring points to constrain contraction into predictable folds^33^. Such control would allow dense neuronal constructs to be shaped to the anatomy of their destination, from aligned tracts for nerve repair to curved sheets matched to a graft site.

The limiting step for translation is throughput. Here we relied largely on manual pipetting, which is sufficient for i*n vitro* studies and disease modeling but introduces operator-dependent variability in construct size and does not scale. Droplet microfluidics is the established route to monodisperse microgels, but collagen is a demanding material for it. Collagen is viscous and thermoresponsive, so the narrow microfluidic geometries needed to form small droplets are prone to clogging. Our system removes this constraint by separating droplet size from final microgel size. Because cells reduce the cast volume, a given target dimension can be reached from a droplet two- to fivefold larger in diameter, so devices can operate with wider channels. As production scales up, core-shell architectures may also be needed to prevent fusion between neighboring microgels. Microfluidics nevertheless requires oil and surfactants with subsequent washing steps and is not always permitted in the biosafety level required for lentiviral vector work. On the other hand, automated liquid handling and extrusion bioprinting, as demonstrated here, avoid both issues and could scale iN-microgel production to the numbers needed for systematic transplantation studies or drug screening.

In direct reprogramming, the donor cell is usually treated as a resource to be overwritten and valued mainly for its accessibility. We show that it can also build the environment it will later occupy. The benefit runs in both directions: matrix remodeling improves reprogramming, and conversion in turn yields a dense tissue the target cell could not assemble. Since contractility is widespread among somatic cells and direct conversion is not restricted to neurons, this principle should extend well beyond fibroblasts and neurons. This establishes a framework in which cell identity and tissue microenvironment are engineered as an integrated process, providing a route toward cell-dense tissue models in which cellular composition, matrix architecture, and cell identity are established together.

## Materials & Methods

### Cell lines

For this study four adult hDF lines were used, these were obtained from the Parkinson’s Disease Research Clinic at the John van Geest Centre for Brain Repair (Cambridge, UK) under full local ethical approval: REC 09/H0311/88 (University of Cambridge). For biopsy sampling, the subjects’ consent was obtained according to the declaration of Helsinki. Details on the biopsy sampling can be found in ^21^. NPCs were generated from the human embryonic stem cell line RC17 using a previously published protocol^67^.

### Cell culture

The hDFs were cultured in uncoated flasks in fibroblast medium consisting of DMEM, 10% fetal bovine serum (FBS), 1% penicillin-streptomycin at 37°C and 5% CO_2_. All media was exchanged every 2-3 days. Upon cells reaching 90% confluency, the cells were split with 0.05% EDTA-Trypsin in DPBS and expanded or used for reprogramming experiments.

Collagen iN-microgels were formed by mixing ice cold 1M HEPES buffer (10% of final volume) with NaHCO_3_ (10% of final volume) and fibroblast media (volume adjusted based on volume collagen, virus and cells needed). The desired volume of rat collagen (R&D Systems) to reach final concentrations of 0.5-3 mg/mL was added to the ice cold mixture to prevent premature polymerization. Next, lentiviral vector cocktails were added to the collagen mixture, for general neuronal reprogramming a polycistronic vector containing *Brn2*, *Ascl1* and 2 short hairpin RNAs (shRNA) targeting *REST* was used at MOI 20. For DA reprogramming, 8 single lentiviral vectors overexpressing *Ascl1*, *Nurr1*, *Lmx1A*, *Lmx1B*, *FoxA2*, *Otx2* and 2 shRNAs for *REST* were used at MOI 5 per vector. For interneuron reprogramming, 5 single vectors overexpressing *Ascl1*, *DLX5*, *LHX6* and 2 shRNAs for *REST* were used at MOI 5 per vector. Lastly, hDFs were added to the mixture (18,000-40,000 cells/microgel depending on intended application). The collagen mixture was then pipetted as droplets onto nontreated plastic and incubated for 25 min at 37°C. Fibroblast media was added on top of the microgel droplets, and a cell scraper was used to gently lift off the droplets from the plastic. Microgels were transferred to ultra-low round bottom plates for further culturing. The day after microgel formation, 80% of the media was replaced with fresh fibroblast media. On day 3 after microgel formation, media was exchanged from fibroblast media to early neuronal reprogramming media consisting of NDiff 227 supplemented with 1% penicillin-streptomycin, CHIR99021 (2 μM), Noggin (50 ng/mL), SB431542 (10 μM), LDN1931189 (0.5 μM), VPA (1 mM), LM22A4 (2 μM), GDNF (2 ng/mL), NT3 (10 ng/mL) and db-cAMP (0.5 mM). Every 2-3 days, 80% of the media was exchanged. On day 16, the media was exchanged to late neuronal reprogramming media consisting of NDiff 227 supplemented with 1% penicillin-streptomycin, LM22A4 (2 μM), GDNF (2 ng/mL), NT3 (10 ng/mL) and db-cAMP (0.5 mM).

For experiments with additional ECM components fibronectin (100 μg/mL), hyaluronic acid (1 mg/mL) and Matrigel (approximately 2 mg/mL) were added together with collagen before the addition of virus and hDFs. iN-microspheres were made as previously published^29^, and the number of cells per sphere was matched with that of iN-microgels.

### Lentivirus production

For reprogramming, third generation lentiviral vectors were produced in HEK-293T cells as previously described in ^68^. In brief, cells were transfected with the plasmid of interest and three helper packing plasmids using transfection agent polyethylenimine (PEI). After 45 h, the virus containing media was filtered with a 0.45 μm filter, ultracentrifuged and resuspended in DPBS. The viruses were stored at –80 °C until use. To determine the titer, RT-qPCR was performed and expression of the virus specific gene (WPRE) was compared against a reference virus with titer determined by FACS.

### Immunocytochemistry and confocal fluorescence imaging

iN-microgels were washed with DPBS twice and fixed with 4% paraformaldehyde (PFA) for 30 min in room temperature (RT). After fixation microgels were washed three times before being stained. Fixed samples were incubated in blocking solution (0.3% Triton X-100, 5% serum and 0.01% sodium azide in PBS) at RT for 24 h. Primary antibodies were diluted in blocking solution (**Table S1**) and incubated for 48-72 h in RT. Following incubation with primary antibodies, cells were washed three times in PBS. Secondary antibodies were diluted in blocking solution (1:200, Jackson ImmunoResearch Laboratories) and DAPI was added at (1:1000). Microgels were incubated with secondary antibody solution for 24-48 h in RT. Lastly, microgels were washed twice and stored at 4°C until imaging.

Microgels were imaged with a NikonA1RHD confocal microscope and a CV8000 (Yokogawa) spinning disk confocal. Microgels were cleared to improve image quality. The microgels were put for 10 min each in 20, 40, 60, 80, and 100% methanol. Next, they were kept in a 2:1 mixture of dichloromethane (DCM) and methanol for 1 hour, followed by two 10 min incubations in 100% DCM. Lastly, the microgels were cleared with ethyl cinnamate and transferred to 96-well black μ-plates (ibidi, #89626) for imaging.

### Scanning electron microscopy

The samples were fixed in 2% glutaraldehyde overnight and rinsed with a 0.1% HEPES buffer. Samples were postfixed for 1 hour in 2% osmium tetroxide before being dehydrated in an ethanol dilution series. This was followed by a dilution series of hexamethyldisilazane (HMDS) before being left to dry in a fume hood overnight. Dehydrated samples were then mounted on 25 mm aluminum stubs, sputter coated with 10 nm platinum/palladium (80:20) and imaged in a JeolJSM-7800F scanning electron microscope.

### Oxygen tension measurements

The technical aspects of the oxygen sensor method, along with sensor integration into cell laden hydrogels, microscopy read-out, and signal processing have been described in detail previously^69,70^. Briefly, oxygen sensor microbeads (CPOx Beads Red, Colibri Photonics GmbH) were embedded in the microgels and microspheres by mixing beads with the cellular suspension at a concentration of 0.7 mg/mL. Oxygen levels were measured through oxygen-mediated quenching of phosphorescence readout with a confocal phosphorescence lifetime microscope (DCS 120 Flim System, Becker & Hickl GmbH). The image processing script was developed and executed in Matlab (Mathworks).

### Numerical modeling of the oxygen distribution

Oxygen diffusion in 96-well ultra-low attachment (ULA) plates seeded with a single 350 µm diameter microsphere or a single 650 µm diameter microgel at its bottom and covered by 100 µL medium was numerically modeled using COMSOL 6.4 (COMSOL AB). Each microsphere or microgel contained 20000 cells evenly distributed across the spherical volume. The oxygen consumption rate was modeled by a Michael-Menten dependence with a Michaelis constant *K_M_* = 6 μM and a maximum consumption rate of *v_max_* = 1.0×10^−17^ mol cell^−1^ s^−1^, as previously measured for hDF cells in 3D cultures^71^. The oxygen diffusion constant was set to *D_O2_* = 3.0 × 10^−9^ m^2^ s^−1^ in culture medium at 37 ℃, and a boundary condition of constant dissolved oxygen concentration at the medium/air interface of *c(O_2_)* = 200 μM (corresponding to gas mixture of 95% air / 5% CO_2_ / 100% relative humidity at 37 ℃) was applied.

### Reflectance microscopy

Samples were scanned at 647 nm excitation on a NikonA1RHD confocal with a 100x Apo TIRF NA 1.49 objective. A nonpolarizing 80:20 beamsplitter was used to collect the reflected light from the sample. Images were processed in ImageJ (NIH).

### Norbornene-modified collagen synthesis

Rat tail collagen type I (Corning,354236) was functionalized with 5-norbornene-2-acetic acid succinimidyl ester (nb-NHS) (Sigma–Aldrich, 776173) at a ratio of 0.1g nb-NHS:1g collagen, as previously described.^72^ Briefly, rat tail collagen was neutralized with NaOH to pH7.2–7.5 and buffered with 10X DPBS to a concentration of 2 mg/mL. Next, nb-NHS was dissolved in dimethylsulfoxide (DMSO) to an initial concentration of 2 mg/ml and diluted 10-fold in 1x PBS. Then nb-NHS solution was added to the neutralized collagen, resulting in a 1 mg/ml final collagen concentration. The reaction proceeded at 4°C for 5 h under rigorous stirring to delay collagen gelation, after which 0.1 N acetic acid was added to quench the reaction and reacidify the collagen solution. The resulting norbornene-modified collagen (nb-collagen) was dialyzed in 25 Mm acetic acid for 4 days, then filtered with a 0.45 μm filter and lyophilized for future use.

### Nb-collagen hydrogel preparation and hDF encapsulation

Lyophilized nb-collagen was dissolved in 0.025 mM acetic acid to a concentration of 8 mg/mL in a low-bind tube, the solution was vortexed on day 0 and 1 and stored for a minimum of 48 h at 4°C until fully dissolved. For nb-collagen hydrogel preparation, ice cold 10X DPBS (10% of final volume) was added to a pre-cooled low bind Eppendorf tube. Next, 1M NaOH (0.022 * volume of needed nb-collagen solution) was added. This was followed by fibroblast media (volume adjusted based on volume of virus and cell suspension needed), lentiviral vectors, and cell suspension. Finally, the ice-cold nb-collagen solution was added to reach a final concentration of 2 mg/mL and mixed carefully. The same procedure as previously described for normal collagen microgels were used for casting and culturing. After 24 h, half of the nb-collagen microgels was treated with the crosslinker methyltetrazine-PEG5-methyhyltetrazine. The crosslinker was dissolved in DMSO to a concentration of 50 mg/mL and then further diluted in fibroblast media to a final concentration of 316 uM. All media was carefully removed from microgels and replaced with the media containing crosslinker before incubation at 37°C for 2 h. Next, microgels were washed once with DPBS before adding fresh fibroblast media for continued culture.

### RT-qPCR

Microgels were treated with 5 mg/mL collagenase in DBPS with 5% FBS for 20 min at 37°C before being washed twice with DPBS. Microgels were then lysed, and RNA was extracted using the RNeasy Micro Kit (Qiagen) following the manufacturer’s instructions. RNA (500 ng – 1 μg) was reverse transcribed to cDNA using the Maxima First Strand cDNA Synthesis Kit (Thermo Fisher). Using the Bravo instrument (Agilent), cDNA, primers of interest (**Table S2**) and SYBR Green Master mix was mixed and analyzed with the LightCycler 480 II instrument (Roche). A two-step protocol with a 30 second denaturation step at 95 °C followed by a 60 second 60°C annealing/elongation step was run for a total of 40 cycles. Samples were normalized against two housekeeping genes, *ACTB* (actin-beta) and *GAPDH* (glyceraldehyde-3-phosphate dehydrogenase) and analyzed using the ΔΔCt-method. The expression was compared to non-converted hDFs and presented as a fold change. Statistical analysis and visualization were performed in GraphPad (Prism 10).

### Bulk RNA sequencing

RNA for Bulk RNA sequencing was extracted using the RNeasy Mini Kit (Qiagen) with DNase treatment according to the manufacturer’s instructions. cDNA libraries were prepared using the Illumina TruSeq library preparation kit and sequenced with 2 x 150 bp paired-end reads on an Illumina NextSeq 2000 using the P3 flow cell giving ~37M reads per sample. Raw base calls were demultiplexed and converted into sample-specific fastq files using bcl2fastq (v2.20, Illumina). Quality control was performed using the nf-core/rnaseq pipeline (FastQC, Trim Galore, STAR, RSeQC and Qualimap) summarized with MultiQC, and all samples were deemed of sufficient quality for downstream analysis. Transcript abundances were quantified with Salmon (v1.9.0) in selective-alignment mode with GC-bias correction against a GENCODE human transcriptome index, and summarized to gene level with tximport using the Ensembl GRCh38 release 99 annotation.

Gene-level counts and transcripts per million (TPM) values were analyzed in R. Gene symbols were annotated using the Ensembl GRCh38 release 99 annotations. For differential expression, genes were retained if they had at least 10 total counts across the samples included in that comparison. Counts were rounded to integers and analyzed using DESeq2. Differential expression was assessed using Wald tests with Benjamini–Hochberg correction for multiple testing, and log2 fold changes were shrunk using apeglm. Genes with adjusted P < 0.05 were considered significant. MA plots were generated in ggplot2 using log10-transformed mean normalized counts (log10(baseMean + 1)) on the x-axis and apeglm-shrunken log2 fold changes on the y-axis.

Principal component analysis (PCA) incorporating fibroblasts and collagen and sphere cultures at day 2, day 7 and day 21 used the 2,000 most variable genes following log2(TPM + 1) transformation. PCAs were performed using centered expression values without scaling genes to unit variance. Curated fibroblast and neuronal marker heatmaps were generated using pheatmap from log2(TPM + 1)-transformed values. Rows and columns were displayed in a predefined order without clustering.

Gene Ontology (GO) Biological Process over-representation analysis was performed using clusterProfiler and org.Hs.eg.db with Ensembl gene identifiers. Genes higher in sphere or collagen cultures at day 21 were analyzed separately, using adjusted P < 0.05 and an absolute unshrunken log2 fold change ≥ 1 to define the input gene sets. Genes with non-missing DESeq2 P values constituted the background. Enrichment analyses used Benjamini–Hochberg correction with P-value and q-value cutoffs of 0.05. Redundant GO terms were simplified using a semantic similarity cutoff of 0.7, retaining the term with the lowest adjusted P value. Fibroblast samples originated from different sequencing runs and comparisons involving fibroblasts are therefore considered exploratory because condition and sequencing-run effects could not be separated.

### Animals

All animal procedures were done according to the guidelines and recommendations of Swedish animal protection legislation and were approved by the local Animal Ethics board (ethical permit no. 5.8.18 00257/2022). Experiments were carried out in adult female athymic rats (Envigo) to avoid graft rejection. All rats were housed in ventilated cages under 12-hour light/dark cycles with free access to food and water. The rats weighed at least 225 grams before the start of the experiment, and weights were recorded weekly to track potential weight loss.

### Surgeries

The surgeries were carried out under general anesthesia given via intraperitoneal injection of Ketaminol (45 mg/kg) and Domitor (0.3 mg/kg). Local anesthesia Marcain (0.1 mL) was given subcutaneously. Rats were places in a stereotaxic frame in a “flat head” position (vertical difference between lambda and bregma < ± 0.2 mm). To ensure widespread depletion of DA neuron in one hemisphere, a unilateral lesion with 6-OHDA (3.5 μg/μL, 3 μL, 0.3 μL/min) in the medial forebrain bundle (MFB) was performed using the following coordinates: A/P: −3.9, M/L: −1.2, D/V: −7.3. DA iN-microgels were transplanted 4 weeks after 6-OHDA lesions in the DA neuron depleted hemisphere with one 3 µL deposit at the injection rate of 1 μL/min using following coordinates A/P: +1.2, M/L: −2.8, D/V: −4.5 (striatum). A 2 min diffusion time was used before retracting the needle. Post-surgery a subcutaneous dose of Antisedan (0.28 mg/kg) and Temgesic (0.04 mg/kg) was given to reverse the anesthesia and serve as an analgesic.

### Preparation of iN-microgels for transplantation

On the day of transplantation, iN-microgels reprogrammed *in vitro* for 7 days were harvested by gentle pipetting through a cut 1 mL tip. iN-microgels were then washed and suspended in Hanks balanced salt solution (HBSS) with 10% DNAse and put on ice until used. Approximately 500,000 hDFs were reprogrammed for transplantation in each animal. To test whether removal of collagen promotes graft-host interaction post-transplantation, iN-microgels were exposed to 5 mg/mL collagenase for 30 min at 37°C before being washed and resuspended in transplantation media.

### Immunohistochemistry

Four weeks after transplantation rats were administered a lethal dose of anesthesia by intraperitoneal injection of sodium pentobarbital. A transcardial perfusion was then performed with room temperature 0.9% saline for 3-5 min followed by ice-cold 4% PFA (pH 7.4 ± 0.2) for 5 min. Following perfusion, brains were manually removed from the skull and left in 4% PFA at 4°C overnight. The next day, PFA was removed and replaced with a 25% sucrose solution. Once the brains had sunk in the sucrose solution (2-3 days), the brains were sectioned on a freezing microtome at 35 μm thickness in 1:8 series and stored in antifreeze at 4°C until staining was started.

Brain sections were stained free-floating in glass vials on a shaker. Sections were washed three times with 0.1M KPBS before performing antigen retrieval by incubating with TRIS-EDTA (pH 9.0) for 30 min at 80°C. Following antigen retrieval, sections were washed three times in KPBS. Sections for DAB staining were incubated in quenching solution (10% H_2_O_2_ and 10% methanol in KPBS) for 15 min at RT before being washed three times with KPBS. Next, sections were incubated with blocking solution (5% serum and 0.25% Trition-X100) for 1 hour at RT. Primary antibodies (**Table S1**) were then diluted in blocking solution and incubated overnight at RT on a shaker. The following day, sections were washed twice in KPBS before being incubated for 30 min in blocking solution. Immunofluorescence staining sections were incubated with fluorophore-conjugated secondary antibodies (Jackson Laboratory 1:200 and DAPI 1:1000) for 1 hour at RT and washed three times with KPBS before being mounted on chrome alum-gelatin coated slide and coverslipped with PVA-DABCO. For DAB staining, sections were incubated with secondary antibody (biotinylated antibody) for 1 hour at RT. Following incubation with secondary antibody, sections were washed three times with KPBS, then incubated in an avidin-biotin complex (ABC Complex) for 1 hour at RT followed by three washes with KPBS. Sections were then incubated with 0.05% DAB-solution for 2 min. Then 20 μL of 3% H_2_O_2_ was added to develop color. Lastly, sections were washed three times with KPBS, mounted on chrome alum-gelatin coated slides and dehydrated in 70%, 95% and 100% ethanol followed by xylene before being coverslipped with DPX mounting media.

### Alginate granular gel preparation

Granular gels are viscoelastic novel materials used for embedded 3D printing as support baths providing high-resolution and enhanced fidelity printing. The granular gel protocol was adapted from two published protocols^70, 73^. Shortly, alginate solution at a concentration of 0.75 wt% was prepared by dissolving 7.5 g of sodium alginate (Fujifilm Wako Pure Chemical Corporation, 196-13325) in 1 L of MilliQ water on a stirring hotplate at 60°C. Once dissolved, it was sterilized through a 0.45 μm pore size filter. Sterile alginate granular gel was prepared by mixing 85 ml of 0.75 wt% alginate solution, 32 ml of pre-sonicated 60mM CaCO3(Merck, C5929) in MilliQ water suspension and 33 ml of 1 wt% Xanthan Gum (Merck, G1253) in a final 150 mM saline solution (Merck, 57653) and stirred for 5 mins at 250 rpm to ensure homogeneous mixing. Gluconic acid solution was generated by dissolving 1.364 g of D-(+)-Gluconic acid δ-lactone (Merck, G4750) powder in 5 ml of MilliQ water and sterilized by passing it through a 0.2 μm pore size filter (Fisherbrand PES, 0.2 μm). The gluconic acid solution was added to the gel mixture to initiate slow release of calcium cations and facilitate uniform internal crosslinking towards a bulk gel. The mixture was then stirred at a lower speed of 100 rpm until the whirlpool disappeared. Then, the gel was set aside to rest overnight at room temperature. The next day, 20 ml of the bulk gel was transferred to a 100 ml blue cap glass bottle containing 80 ml of MilliQ water and homogenized at 25000 rpm for 2 min using a T25 digital Ultra-Turrax disperser control (IKA) with an appropriate dispersing head tool (IKA, S25N-10G-ST) for 100 mL volume. The homogenate was divided evenly into two 50 mL falcon tubes. The whole process was repeated until the whole bulk gel was homogenized. The falcon tubes were incubated at 4°C for 24 h, followed by centrifugation at 18500 g and removal of the supernatant. The recovered gel pellets were washed with 20 ml DMEM/F-12 (Gibco, 10565018) media, vortexed vigorously and stored at 4°C. Before printing, the falcon tubes were centrifuged at the same parameters as before so the gel pellet could be recovered.

### Embedded 3D printing

Embedded 3D printing was employed in this study to generate fibroblast-collagen structures with well-defined 3D geometry. Cell ink was prepared and loaded into a 250 μl glass gastight syringe (Hamilton, #1725) with a 22G and 0.5″ length needle (CELLINK). Embedded 3D printing was performed using a BioAssemblyBot 400 (Advanced Solutions) with integrated 3D CAD software for designing structures and that automatically generate the toolpath. The printed cylinder comprised of six homocentric 8 mm diameter circles with 400 μm spacing in the z-direction. The loaded syringe was placed into a mechanical head tool while the lateral speed and extrusion rate were set at 0.5 mm/sec and 20 ul/sec respectively. Once the structures were printed in a middle of a 24 well plate (Corning, CLS3527) they were transferred to an incubator to facilitate the crosslinking of the collagen.

### Nanoindentation

Nanoindentation measures local stiffness (Young’s modulus) from the load-displacement response of a probe indenting the sample surface. Here, we used a Pavone Nanoindenter (Optics11 Life) fitted with a probe with cantilever stiffness of 0.47 N/m, tip radius of 26 μm and geo factor in air of 2.76. Before each experiment, the probe was calibrated according to manufacturer guidelines in the same media used for cell culturing. Stiffness experiments consisted of two steps: finding the surface by probe moving at 50 µm/s speed and retracting 20 µm from it, followed by a two-point matrix scan along the Y-direction with 5 µm spacing and a 20 µm Z-retraction between points. Peak load poking profile with max load of 0.1 µN and piezo speed of 5 µm/s was selected as the indentation mode of operation. All measurements were performed approximately at the center of each sample.

### Statistical information

Statistical analysis and plotting were performed in GraphPad Prism. p values are marked as following in figure panels: *, P < 0.05; **, P < 0.005; ***, P < 0.0005; ****, P < 0.00005. Details on statistical analysis of the bulk RNA-seq data is described in the corresponding materials and methods section. Statistical analysis for the rest of the figure panels was performed as follows: Fig. 1e – unpaired one-way ANOVA (F = 397.8, DF = 11) Fig. 1g – unpaired t-test (t = 11.97, DF = 6); Fig. 3f – one-way ANOVA (MAP2: F = 2.08, DF = 11; MAPT: F = 2.24, DF = 7; SYN1: F = 1.46, DF = 11; SYP: F = 0.86, DF = 11; NCAM: F = 0.48, DF = 11; COL1A1: F = 1.64, DF = 11; VIM: F = 1.07, DF = 11); Fig. 4a – one-way ANOVA (F = 8.18, DF = 49); Fig. 4b – one-way ANOVA (MAP2: F = 7.68, DF = 11; MAPT: F = 0.67, DF = 11; SYN1: F = 13.62, DF = 11; SYP: F = 4.47, DF = 11; NCAM: F = 1.64, DF = 11; COL1A1: F = 1.14, DF = 11; VIM: F = 0.22, DF = 11); Fig. 4c – one-way ANOVA (MAP2: F = 3.3 DF = 11; COL1A1: F = 1.08, DF = 11); Fig. 4f – unpaired t-test (t = 5.42, DF = 22.2); Fig. 4g – unpaired t-test (t = 1.03, DF = 3.98); Fig. S2 – one-way ANOVA (MAP2: F = 21.05, DF = 11; MAPT: F = 6.65, DF = 7; SYN1: F = 14.32, DF = 11; SYP: F = 12.28, DF = 11; NCAM: F = 0.48, DF = 11; COL1A1: F = 14.58, DF = 11; VIM: F = 1.07, DF = 11).

## Acknowledgments

The authors would like to thank Michael Sparrenius and Sara Corsi for the help with animal work; Bengt Mattsson for providing help with brain slide imaging; Sol da Rocha Baez for the help with virus production; Petter Storm for the help with bioinformatics analysis; Jonas Viswalingam Bagge (StemScreen, University of Copenhagen, IFD grant No. 2078-00002B) for the help with automated confocal fluorescence imaging and image analysis. Lund University Bioimaging Centre (LBIC) is gratefully acknowledged for providing experimental resources for confocal reflectance and electron microscopy. We would also like to thank Ross Cullen Bretherton and Charis Segeritz-Walko from Novo Nordisk A/S for providing access to and training for the nanoindentation tool.

This work was supported by funding from the New York Stem Cell Foundation (MP), European Research Council (ERC) under ERC Grant Agreement 771427 (MP), European Union–funded project NSC-Reconstruct (European Union, H2020, GA no 874758, 2020-23, MP), Swedish Research Council (MP 2021-00661, MP 2021-02967, and DRO 2021-01839), Swedish Parkinson Foundation (Parkinsonfonden, MP), Swedish Brain Foundation (MP), Konung Gustaf V:s och Drottning Victorias Frimurarestiftelse (MP), Olle Engkvist Foundation (213-0229, DRO), the Lundbeck Foundation (R380-2021-1267, JK). The Novo Nordisk Foundation Center for Stem Cell Medicine is supported by a Novo Nordisk Foundation grant (NNF21CC0073729). The work was also supported by the National Institute for Health Research Cambridge Biomedical Research Centre, (NIHR203312). The views expressed are those of the authors and not necessarily those of the NIHR or the Department of Health and Social Care. The funding bodies played no role in the design of the study or in collection, analysis, or interpretation of data or in writing of the manuscript.

## Contributions

Conceptualization: JK, KL, FN, MP

Cells, reagents, analytical tools: RAB, DRO, MFW, NBL, EBR, DJM

Investigation and analysis: KL, JK, FN, PA, JG, AB

Bioinformatics analysis: MH

Writing - original draft: KL, JK

Writing - review & editing: KL, JK, MH, MP, RAB, JG, AB, DRO

## Data Availability

The bulk RNA-seq data generated in this study have been deposited in the GEO database under accession code GSE351887 (https://www.ncbi.nlm.nih.gov/geo/query/acc.cgi?acc=GSE351887). This includes gene-level read counts and TPM values for all samples. The fibroblast and 3D iN-microsphere samples reanalyzed in this study were previously published and are available in the GEO database under accession code GSE287155 (samples GSM8740065–GSM8740067 and GSM8740080–GSM8740088). All other data supporting the findings of this study are available from the corresponding author upon request.

