## Supplementary material for "Transient matrix remodeling during direct cell-fate reprogramming enables self-assembly of dense neuronal niches"

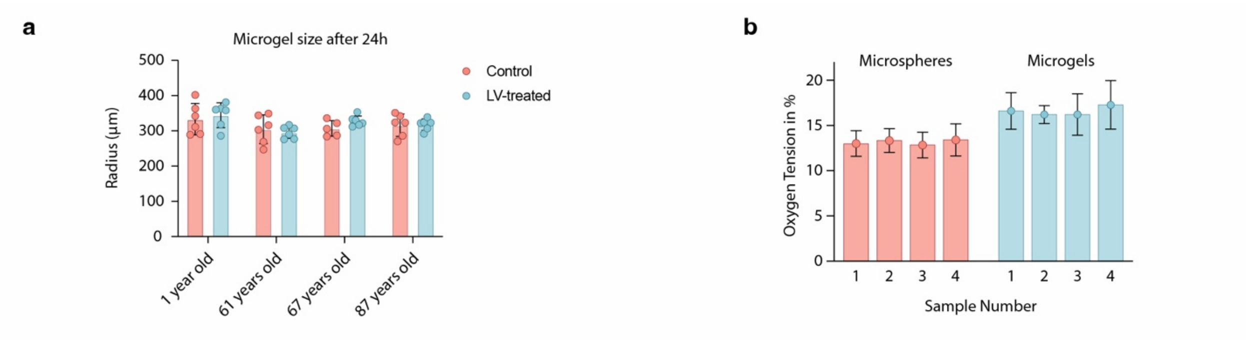


**Fig. S1 a** Measurements of microgel radii 24 hours after gel casting, with and without lentivirus (LV) added, for four hDF lines. **b** Oxygen tension measurements showing mean ± s.d. for each measured microsphere or microgel.

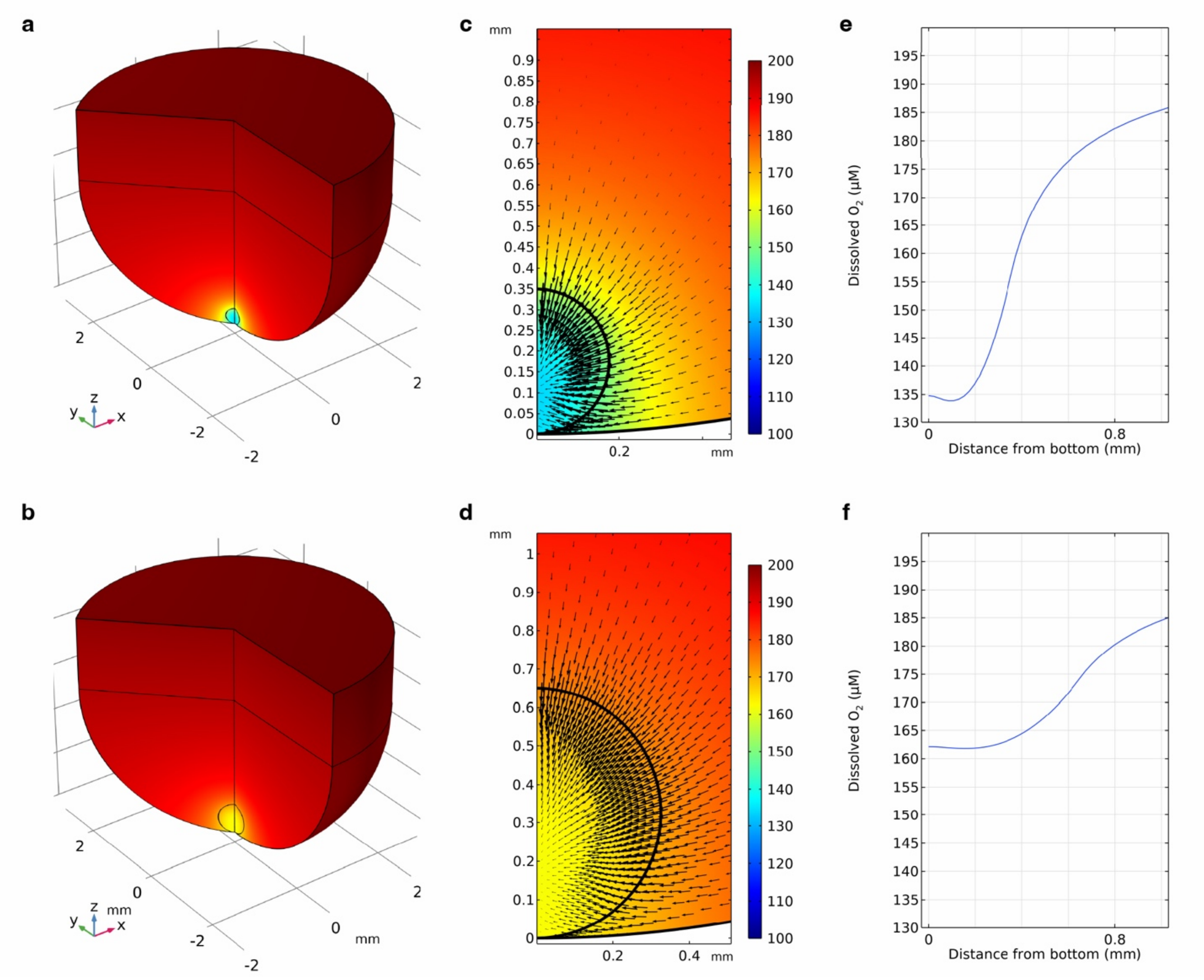


**Fig. S2** Steady-state dissolved oxygen concentration in ULA wells containing a 350 μm diameter microsphere (**a** sector cut-away of the culture medium) or a 650 μm diameter microgel (**b**), each consisting of 20000 hDF cells. The lower surface-to-volume ratio of the microsphere (**c**; cross-sectional view of the microsphere and proximal culture medium) compared to the microsphere (**d**) limits the influx of oxygen, resulting in lower oxygen concentrations in the microsphere (color legend in μM dissolved oxygen applies to sub-figures **a**-**d**). Vertical oxygen concentration profiles from the well bottom through the microsphere (**e**) and the microgel (**f**).

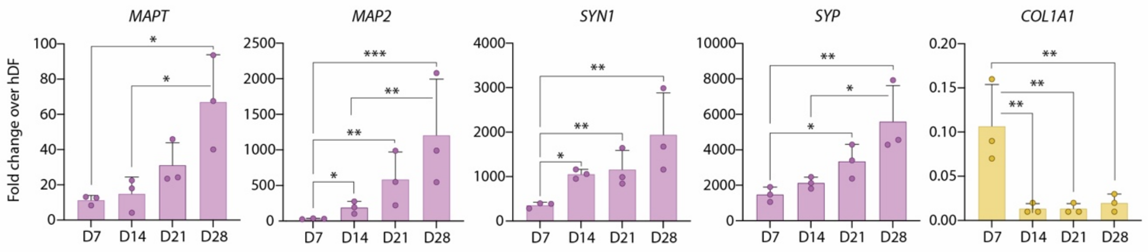


**Fig. S3** RT-qPCR analysis of selected fibroblast and neuronal genes of iN-microgels at day 7, 14, 21 and 28 with each dot representing different hDF line. * p <0.05, ** p < 0.005, *** p < 0.0005

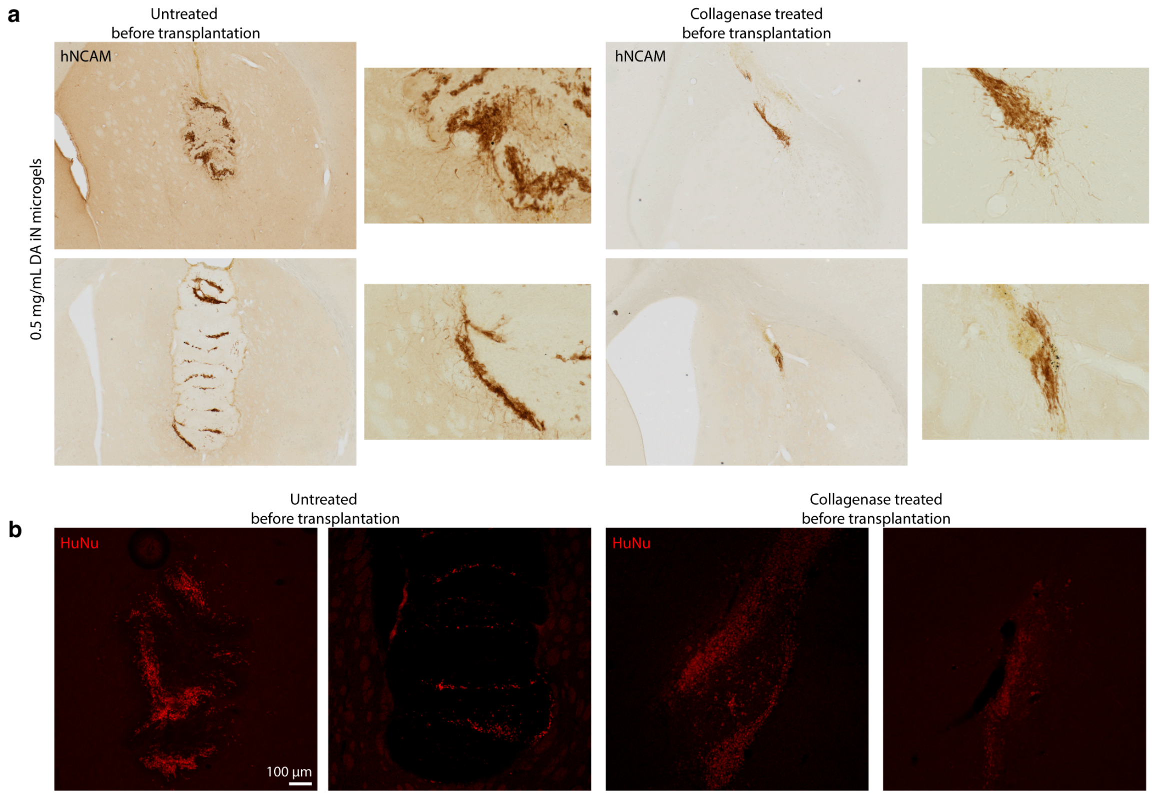


**Fig. S4** Transplantation of 0.5 mg/mL iDAN-microgels in 6-OHDA lesioned nude rats. **a.** DAB-staining of grafted iDAN-microgels 4 weeks post transplantation visualized by human NCAM. Grafts with non-treated microgels to the left and microgels treated with 5 mg/mL collagenase prior to transplantation to the right. Images depict an overview of grafts with zoom in to display fibers. **b.** Immunofluorescent human nuclei (HuNu) staining the grafts of untreated and treated iDAN-microgels.

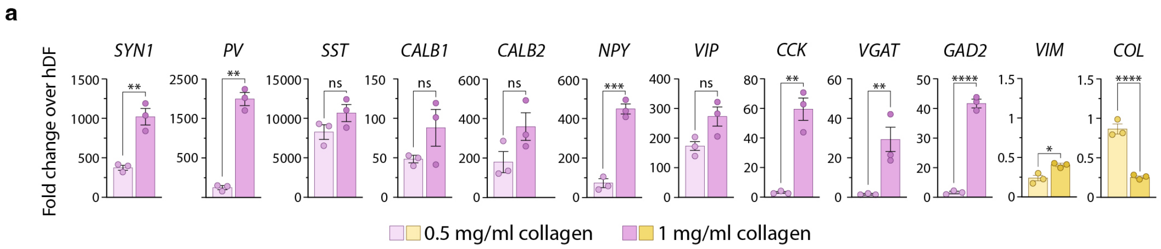


**Fig. S5** RT-qPCR analysis comparing expression of interneuron and fibroblast genes at day 28 in induced GABAergic interneuron microgels using two collagen concentrations, 0.5 and 1 mg/mL.

**Table 1 - List of primers used for RT-qPCR analysis of reprogrammed iN-microgels.**

| Primers | | Sequence (5'-3') | Full gene name |
|---|---|---|---|
| *AADC* | F | GGGGACCACAACATGCTGCTCC | DOPA decarboxylase (DDC) |
| | R | AATGCACTGCCTGCGTAGGCTG | |
| *ACTB* | F | CCTTGCACATGCCGGAG | Beta-actin |
| | R | GCACAGAGCCTCGCCTT | |
| *CALB1* | F | TGGCTCCATTTCGACGCTGACG | Calbindin |
| | R | ATCCAGCCTTCTTTCGCGCCTG | |
| *CALB2* | F | TGGAGGCTTGGCGGAAGTACG | Calretinin |
| | R | CCGGTTCGCCTTCTTCAGCAG | |
| *COL1A1* | F | TGAAAATGGAGCTCCTGGTCAG | Collagen type 1 alpha 1 |
| | R | AGCACCAGTAGCACCATCATTT | |
| *DAT* | F | CACTGCAACAACTCCTGGAA | Dopamine transporter (SLC6A4) |
| | R | AAGTACTCGGCAGCAGGTGT | |
| *GAD2 (GAD65)* | F | ATCCTCACGACTCAGCTCCC | Glutamate decarboxylase 2 |
| | R | GAGCTTTAAAAGAGACCGGGACT | |
| *GAPDH* | F | TTGAGGTCAATGAAGGGGTC | Glyceraldehyde-3-phosphate dehydrogenase |
| | R | GAAGGTGAAGGTCGGAGTCA | |
| *MAP2* | F | CCGTGTGGACCATGGGGCTG | Microtubule associated protein 2 |
| | R | GTCGTCGGGGTGATGCCACG | |
| *MAPT* | F | TGTGGGTCAGTGTGCCACCCT | Microtubule associated protein Tau |
| | R | CACGAAGTGCCACCCTCCTGC | |
| *NCAM* | F | GTCAGAGGCCACCGTCAACGT | Neural adhesion molecule 1 |
| | R | CTTCCCCCTCCCGGAACTCCT | |
| *PITX3* | F | GGAGGTGTACCCCGGCTACTCG | Paired-like homeodomain 3 |
| | R | GAAGCCAGAGGCCCCACGTTGA | |
| *PVALB* | F | TGCAGGATGTCGATGACAGA | Plasmalemma vesicle-associated protein |
| | R | TTTCTTCAGGCCGACCATTT | |
| *SST* | F | CAAGCCGCTTTAGGAGCGAG | Somatostatin |
| | R | ACGCGGCAGGACAGCATCT | |
| *SYN1* | F | CCCGTGGTTGTGAAGATGGGGC | Synapsin 1 |
| | R | TGCCACGACACTTGCGATGTCC | |
| *SYP* | F | ACCTCGGGACTCAACACCTCGG | Synaptophysin |
| | R | GAACCACAGGTTGCCGACCCAG | |
| *TH* | F | CGGGCTTCTCGGACCAGGTGTA | Tyrosine hydroxylase |
| | R | CTCCTCGGCGGTGTACTCCACA | |
| *VIM* | F | ACAAAAAGTCCGCACATTCGAG | Vimentin |
| | R | ACAAGAGCGCCCCTAAGTTTTT | |

**Table 2 - primary antibodies used in this study.**

| Marker | Specificity | Source (cat. no.) | Dilution |
|---|---|---|---|
| Calbindin | Rabbit | Swant (CB38) | 1:500 |
| Calretinin | Rabbit | Abcam (ab702) | 1:200 |
| COL1A1 | Sheep | R&D Systems (AF6220) | 1:500 |
| GABA | Rabbit | Sigma (A2052) | 1:2,000 |
| GAD65/67 | Rabbit | Abcam (ab5392) | 1:2,000 |
| hNCAM | Mouse | Santa Cruz (sc 106) | 1:500 |
| HuNu | Mouse | Merck Millipore (MAB1281) | 1:200 |
| MAP2 | Chicken | Abcam (ab5392) | 1:5,000 |
| NeuN | Rabbit | Millipore (ABN78) | 1:1,000 |
| Parvalbumin (PV) | Rabbit | Swant (PV27) | 1:500 |
| Somatostatin (SST) | Mouse | Santa Cruz (sc55565) | 1:200 |
| TAU | Mouse | Thermo Fisher (MN1000) | 1:500 |
| TAU | Rabbit | DAKO (A0024) | 1:500 |
| TH | Rabbit | Merck Millipore (AB152) | 1:1,000 |
| TH | Sheep | Millipore (AB1542) | 1:200 |
| TUBB3 | Mouse | BioLegend (801202) | 1:500 |
| vGLUT | Mouse | SYSY (135311) | 1:500 |
